# The effect of sleep continuity disruption on multimodal emotion processing and regulation: a laboratory-based, randomized, controlled experiment in good sleepers

**DOI:** 10.1101/2022.04.22.489209

**Authors:** MJ. Reid, X. Omlin, CA. Espie, R. Sharman, S. Tamm, SD. Kyle

**Affiliations:** Sleep and Circadian Neuroscience Institute, The University of Oxford, Oxford, UK; Johns Hopkins School of Medicine, Baltimore, MD, USA; Department of Psychiatry, The University of Oxford, Oxford, UK; Department of Clinical Neuroscience, Karolinska Institutet, Stockholm, Sweden

**Author notes:** Corresponding Author: Dr Matthew J Reid, Department of Psychiatry and Behavioral Sciences, Johns Hopkins School of Medicine, 5510 Nathan Shock Drive, Baltimore MD. **Disclosure Statement:** Financial Disclosure: none. Non-financial Disclosure: none This research study was supported financially by the National Institute for Health Research (NIHR) Oxford Biomedical Research Centre (BRC). The research was also supported through a DPhil (PhD) Scholarship (to M.J.R) from the Dr Mortimer and Theresa Sackler Foundation. **Author Contributions:** M.J.R designed, conducted the research and wrote the manuscript, X.O assisted in collection of the data and study experiment procedures, C.A.E and S.D.K supervised the study design and conduct, and assisted in writing the manuscript. S.T assisted in writing and editing the manuscript, R.S assisted in data collection and EEG scoring. All authors provided feedback and contributed to the final version.

**Keywords:** Sleep Deprivation, Depression, Attentional Bias, Memory Consolidation, Emotional Regulation, Sleep, Emotion

## Abstract

Previous research shows that experimental sleep deprivation alters emotion processing, suggesting a potential mechanism linking sleep disruption to mental ill-health. Extending previous work, we experimentally disrupted sleep continuity in good sleepers and assessed next-day emotion processing and regulation using tasks with established sensitivity to depression. In a laboratory-based study, 51 good sleepers (37 female; mean age = 24 years, SD= 3.63) were randomized to one night (23:00-07:00) of uninterrupted sleep (n=24) or sleep continuity disruption (n=27). We assessed emotion perception, attention, and memory the following day. Participants also completed an emotion regulation task and measures of self-reported affect, anxiety, sleepiness, overnight declarative memory consolidation, and psychomotor vigilance. Confirming the effects of the manipulation, sleep continuity disruption led to a marked decrease in polysomnography-defined total sleep time (229.98 mins vs 434.57 mins), increased wake-time after sleep onset (260.66 mins vs 23.84 mins) and increased sleepiness (d=0.81). Sleep continuity disruption led to increased anxiety (d=0.68), decreased positive affect (d=-0.62), reduced overnight declarative memory consolidation (d=-1.08) and reduced psychomotor vigilance [longer reaction times (d=0.64) and more lapses (d=0.74)], relative to control. However, contrary to our hypotheses, experimental sleep disruption had no effect on perception of, or bias for, emotional facial expressions, emotional memory for words, or emotion regulation following worry induction. In conclusion, one night of sleep continuity disruption had no appreciable effect on objective measures of emotion processing or emotion regulation in response to worry induction, despite clear effects on memory consolidation, vigilance, and self-reported affect and anxiety.

## Introduction

Previous research indicates a link between sleep and emotional functioning (Baglioni, Spiegelhalder et al. 2010, Krause, Simon et al. 2017). Dysfunctional emotional processing has been suggested as a potential mechanism underlying the association between sleep disruption and psychiatric disorders (Hertenstein, Feige et al. 2019). In support of this proposition, several early studies reported that total sleep deprivation (TSD) negatively affects processing of emotional stimuli, including facial expressions (Van Der Helm, Gujar et al. 2010), as well as memory for emotional words and images (Phelps 2004). However, recent studies have failed to show an effect of sleep deprivation (or restriction) on emotional processing (Holding, Laukka et al. 2017, Gerhardsson, Åkerstedt et al. 2019, Tamm, Schwarz et al. 2020).

TSD and chronic sleep restriction protocols do not adequately reflect the type of sleep disruption experienced by patients with mental health problems, where the chief defining feature is sleep discontinuity and self-reported insomnia. Case-control studies, comparing participants with and without insomnia, have also found behavioral impairments in emotional tasks, such as reduced intensity ratings of emotional faces (Kyle, Beattie et al. 2014), and reduced recognition of positive emotional images (Chunhua, Jiacui et al. 2019). However, similar to the TSD literature, inconsistent findings are observed (Baglioni, Spiegelhalder et al. 2010, Crönlein, Langguth et al. 2016), sample sizes have typically been small, and a broad range of tasks have been used across studies, which vary in both their demands and sensitivity to specific cognitive processes. It is clear that the field needs to causally test the effects of sleep manipulation protocols that map onto sleep disturbance experienced by clinical populations. Moreover, there is a need to investigate different facets of emotional functioning within the same study (using standardized tasks), including processing of emotional stimuli across multiple cognitive domains, regulation of emotion in response to challenge, and self-reported mood state.

We designed a study to address these needs and to assess the effects of sleep continuity disruption on emotional functioning. We selected a forced awakenings protocol (Finan, Quartana et al. 2017) in order to simulate sleep continuity disruption and architectural impairment characteristic of a severe insomnia phenotype (Finan, Quartana et al. 2017). Previous work implementing this protocol has demonstrated attenuation of positive affective systems in healthy participants without influencing negative affect, resulting in an overall negativity bias (Finan, Quartana et al. 2017). Advancing previous research, we focussed on cognitive-affective tasks with established sensitivity to depression risk, diagnosis, and treatment (Harmer, Cowen et al. 2011), and investigated emotion regulation as well as mood state. We hypothesized that sleep continuity disruption would lead to enhanced processing of negative emotional stimuli and reduced processing of positive emotional stimuli in the following domains: 1) recognition of facial emotions, 2) attention towards emotional faces, 3) categorization of emotional words, and 4) memory for emotional words (see supplemental table 3 for detailed hypotheses). We used a worry induction task to assess effects on goal-oriented attention and positive/negative thought intrusions, hypothesizing impairment in the sleep continuity disruption group versus control (see methods for detailed hypotheses). In order to confirm the effects of the sleep manipulation, and to provide context in terms of magnitude of group effects, we assessed affect, sleepiness, vigilance and overnight memory consolidation; domains that are known to be sensitive to the effects of sleep loss.

## Methods

### Design

We performed a between-subjects randomized controlled experiment to assess the effects of sleep continuity disruption (delivered via forced awakenings(Finan, Quartana et al. 2017)) on emotion processing and emotion regulation in a laboratory setting (see figure 1 for schematic of tasks). Our primary outcomes were emotional perception (facial emotion recognition test), emotional attention (faces dot probe task), and emotional memory (emotional recall and recognition memory tasks) measured using the Oxford Emotional Test Battery [ETB (P1Vital Product ltd, Oxford UK)], and emotion regulation measured using an established worry induction (Adams, Pounder et al. 2016). A between-group design was employed because our primary outcome measure tasks may be susceptible to practice effects (Adams, Pounder et al. 2016). The study was given approval from The Medical Sciences Interdivisional Research Ethics Committee (MS IDREC), University of Oxford Central University Research Ethics Committee (CUREC).

**Figure 1.**
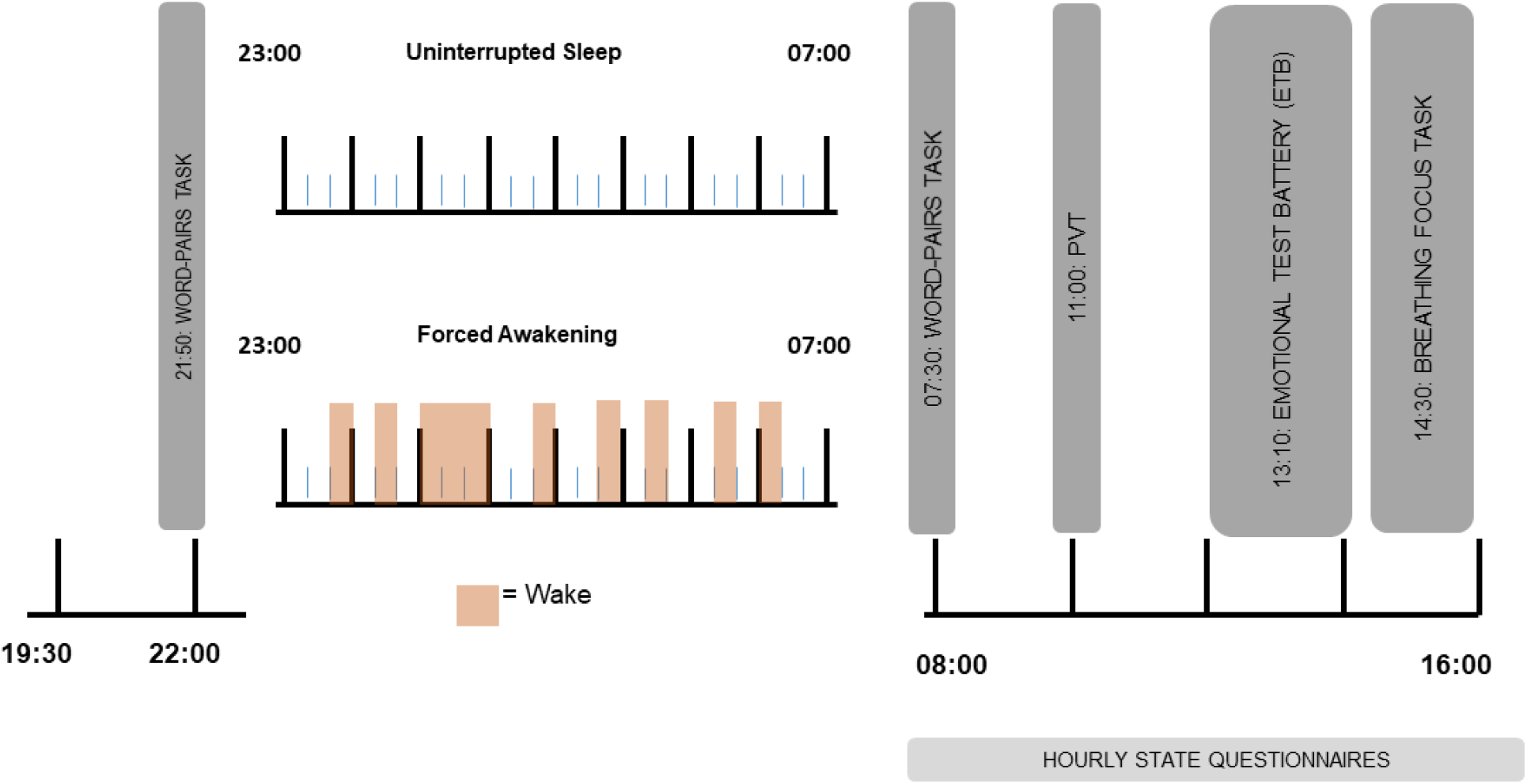
Timing of evening and daytime tasks. Task timings are standardized across participants and tasks were completed a minimum of 30 mins pre and post sleep. The timing of sleep protocols and forced awakening periods were also fixed across participants.

We recruited healthy participants (aged 18-30 years, 38 female; mean age = 24 years, SD= 3.63) who were habitual good sleepers, free from any current or pre-existing sleep, neurological or psychiatric disorders, and were free of any centrally acting central nervous system medications (see inclusion criteria in table 1).

**Table 1 :** Inclusion and exclusion criteria.

| Inclusion | Exclusion |
| --- | --- |
| • 18-30 years<br>• Low caffeine users ($\leq 400\text{mg}/4$ cups of coffee per day)<br>• Non-smoker<br>• Low-moderate drinker ($\leq 2$ drinks/ 4 units daily)<br>• PSQI Total Score $\leq 5$.<br>• PSQI Item 6 = 'Very Good' or 'Fairly Good'<br>• Usual sleep onset latency $\leq 30$ mins, defined by PSQI items (2 & 5a)<br>• Endorsed good habitual sleep quality<br>• MEQ score 42-58.<br>• Usual total sleep time between 7-9 h/ night defined by PSQI measure item 5.<br>• HADS-Depression $\leq 7$ Score & HADS-Anxiety scores $\leq 7$ | • Report being troubled by sleep<br>• Report current or previous diagnosis of mental health, or substance abuse disorder<br>• Alcohol dependence (AUDIT score $> 8$)<br>• Report current or lifetime history of neurological disorders or traumatic brain injury/ significant loss of consciousness<br>• Report current or lifetime history of sleep disorders: Insomnia, Narcolepsy, Sleep Apnea, Nightmare Disorder, Night Terrors, Circadian Rhythm Disorders, Restless Leg Syndrome<br>• Report current or lifetime history of medical or cardiovascular disorders<br>• Learning difficulties or intellectual disabilities<br>• Report insufficient understanding of English<br>• Reported use of prescription or recreational drugs<br>• Currently engaged in nocturnal or rotating shift-work<br>• Trans-meridian travel $\geq 4$ hours within the past month or planned within the next month<br>• Report of pregnancy, or planned pregnancy within 2 months |

Abbreviations: MEQ = Morningness Eveningness Questionnaire, HADS =Hospital Anxiety and Depression Rating Scale, PSQI = Pittsburgh Sleep Quality Index, AUDIT = Alcohol Use Disorders Identification Test BSDS: Brief Sleep Disorders Screen.

### Procedures

#### Recruitment

Participants were recruited through several channels, including online, print, and email. Online and written consent was obtained prior to any procedures being undertaken. Participants first completed an online screening questionnaire followed by a study screening visit at the University of Oxford to assess eligibility according to inclusion and exclusion criteria. The following questionnaires were used during the screening procedure: Pittsburgh Sleep Quality Index (Buysse, Reynolds et al. 1989) (PSQI), Morningness Eveningness Questionnaire (Adan and Almirall 1991) (MEQ), Hospital Anxiety and Depression Rating Scale (Bjelland, Dahl et al. 2002) (HADS), Alcohol Use Disorders Identification Test (Babor, Higgins-Biddle et al. 2001) (AUDIT) and Brief Sleep Disorders Screen (Espie 2006) (BSDS). Baseline measures of neuroticism (Eysenck Personality Questionnaire – Revised Short Form (EPQ-RS) (Eysenck, Eysenck et al. 1985) and rumination (Ruminative Responses Scale (Treynor, Gonzalez et al. 2003) (RRS)) were also obtained during this phase in order to characterize the sample at baseline. Participants received reimbursement for participation (£100 in total).

#### Sleep monitoring phase

To further determine eligibility, participants underwent an at-home monitoring phase consisting of seven days of actigraphy concurrent with sleep diaries (to index total sleep time [TST] and sleep efficiency [SE] parameters). During this phase, participants were advised to maintain their regular habitual sleep schedule. Participants wore actiwatches (MotionWatch 8, CamnTech Ltd, Cambridge, UK) 24 hours per day on their non-dominant wrist. Data from the night-time periods were used to calculate average TST and SE using validated algorithms (Elbaz, Yauy et al. 2012) embedded in the actigraphy software (MotionWare 1.2.5, CamnTech, Cambridge, UK). Consistent with previous studies, participants indicated bedtime and risetime using the actigraphy event marker, completed the consensus sleep diary (Carney, Harris et al. 2010) upon waking each morning. Diaries were used to verify bed and rise times of actigraphy measures and gather self-report data on perceived TST and SE. In the instance of missing event marker data, sleep diaries were used to substitute bedtime and rise time values in the actigraphy software.

#### Experimental protocol

Immediately after completion of the seven-day screening phase, participants arrived at the sleep laboratory at 19:30. In the daytime hours prior to coming to the lab, we requested participants abstain from all alcohol, caffeine and stimulants, and to not engage in intense exercise. Participants were set up for polysomnographic (PSG) recording (SomnoMedics GmbH, Germany) and subsequently completed the first session of the computerized memory assessment (word-pairs task (Marshall, Helgadottir et al. 2006) consisting of encoding and recall). Participants were randomized at 22:30 and, following completion of the memory task, were informed about which sleep protocol they were to undergo. At this point, both experimenter and participant were necessarily un-blinded. Commencement of sleep opportunity for both groups began at 23:00. Access to smartphones or handheld electronic devices was restricted to before 22:30 on the first night and between the hours of 07:00-07:30 the following morning. For the duration of the protocol (19:30 to 16:00 the next day) participants remained in the laboratory. Questionnaires assessing state affect (Positive and Negative Affective Schedule: PANAS (Crawford and Henry 2004), emotion regulation (State Difficulties in Regulation Scale: S-DERS (Lavender, Tull et al. 2015), anxiety (State-Trait Anxiety Inventory: STAI (Spielberger 1970), and sleepiness (Karolinska Sleepiness Scale: KSS (Kaida, Takahashi et al. 2006)) were collected pre-randomization to compare groups at baseline, and completed hourly the following day commencing at 08:00 (Post sleep disruption). During the time in the sleep lab, participants did not consume any caffeinated drinks. Standardized meals were consumed at fixed times (09:00, 12:00).

PSG was conducted according to the American Academy of Sleep Medicine (AASM) standards (Berry, Brooks et al. 2017). A standardized PSG montage consisting of 8 EEG electrodes (F3, F4, F7, F8, C3, C4, O1, O2), 2 electrooculography (EOG) electrodes (EOG1, EOG2), 2 submental electromyography (EMG) electrodes and 2 electrocardiogram (ECG) electrodes (to facilitate artefact detection) were used to record objective sleep using a SOMNO HD device (SomnoMedics GmbH, Germany) between 23:00 and 07:00. We opted not include further EMG and respiratory channels required to perform sleep apnoea or limb movement disorder screening, to optimise participant comfort, as the probability of detecting occult sleep disorders in our highly selected healthy sample was low. Electrode sites were measured using the international 10-20 EEG system and positioned according to AASM guidelines. EEG channels were recorded with a sampling rate of 256hz (hardware filter: high pass = 0.3hz, low pass = 128.0hz), using Cz as reference and FpZ as the ground electrode. For analysis signals were re-referenced to the contra-lateral mastoid (M1 and M2). PSG records were scored by a trained scorer according to current AASM guidelines (Scoring Manual Version 2.5 (Berry, Brooks et al. 2012)). A concordance check was carried out on 15% of records by a European Sleep Research Society (ESRS) Somnologist, achieving a concordance rate of >90%.

#### Randomization procedures

Participants were randomly allocated 1:1 to the forced awakenings group (experimental) group or the uninterrupted sleep group (control), stratified according to sex to minimise influence on baseline differences in sleep architecture and emotion. Randomisation schedules were created off-site by a member of the research team with no direct involvement in the study using randomly assigned permuted blocks ranging in size between 4-8. Allocations were stored in sealed opaque envelopes until after the pre-sleep tasks and questionnaires had been completed (22:30). When two participants were present in the laboratory on the same night, participants were randomized as a dyad to prevent forced awakenings from disrupting unrestricted sleep conditions. Given the nature of the manipulation, neither participants nor researchers could be blinded. Nevertheless, both participants and researchers were blinded until commencement of the sleep protocol, including during assessments and tasks conducted pre-sleep. Researchers did not have access to future allocation sequences, which was handled by a study member with no participant contact.

#### Sleep manipulations

Between the hours of 23:00-07:00 participants underwent their assigned sleep group whilst ambulatory PSG recordings were obtained. During this time participants in both groups remained in the bedrooms and access to watches, clocks, mobile phones and other handheld electronic devices was restricted. Participants were permitted to leave the room only to use the bathroom.

##### Forced awakening (Experimental)

Participants underwent sleep continuity disruption via a standardized forced awakening protocol (Finan, Quartana et al. 2017). The night (23:00-07:00) was divided into eight 1-hr intervals. According to a previous study procedure one of these intervals is randomly allocated as a 60-minute awakening. The remaining seven 1-h intervals are then subdivided into thirds (20-min intervals), and one 20-min interval was selected within each hour as a forced awakening period. In the present experiment, we purposefully fixed the timing of awakenings and sleep times between participants, and allocated the third interval as a 60-min awakening, to promote interruption of slow wave sleep. This adaptation was made as previous forced awakening studies showed reductions in SWS mediated impairment in positive affect (PANAS +ve scores) observed in participants undergoing sleep continuity disruption (Finan, Quartana et al. 2015, Finan, Quartana et al. 2017). The maximum possible sleep time was 280mins (WASO = 200 mins) divided across eight awakenings. To wake participants up, a researcher entered the room to wake participants and remained in the room to ensure continuity of wakefulness.

##### Uninterrupted sleep (Control)

Participants were given an eight-hour sleep opportunity between 23:00 and 07:00 and instructed to sleep as normal during this period. Participants in the uninterrupted sleep

### Outcome measures

Participants completed a series of assessments in the evening before sleep, and throughout the subsequent daytime period according to a standardized schedule (+/-5 mins). Testing ceased 30 minutes before bed and commenced 30 minutes post rise-time. Emotion processing tasks were selected on the basis of having a prior demonstration of sensitivity to depression(Harmer, Cowen et al. 2011) and to capture key cognitive-emotional processes (emotional perception, attention and memory). The emotion regulation task (Hirsch, Hayes et al. 2009) was selected on the basis of providing an ecologically valid assessment of trans-diagnostic processes (decreased attentional capacity and increased intrusive thoughts) relevant to psychiatric disorder, including depression and insomnia.

### The Oxford Emotional Test Battery (ETB)

The Oxford Emotional Test Battery (ETB) is a cognitive test battery used to provide a multimodal assessment of emotional biases in the domains of memory, attention and perception (Adams, Pounder et al. 2016). Constituent tasks include the facial expression recognition task (FERT), the emotional categorisation task (ECAT), the faces dot-probe task (FDOT), the emotional recall task (EREC), and the emotional recognition memory task (EMEM). The battery takes approximately 60 mins to complete. The battery was administered after lunch, around 13:00 on the day following sleep manipulation, minimizing the risk that the circadian peaks in wakefulness and mood that typically occur mid-morning (McClung 2013) could mask the effects of the sleep disruption. The battery was delivered using a custom computer monitor (at 60 cm distance) and a button box (P1Vital Products Ltd). For detailed methods of the constituent tasks see supplement.

### Worry induction challenge

A worry induction task was conducted to provide an ecologically valid assessment of emotion regulation in response to challenge (a worry induction). The worry induction task procedure followed a protocol derived from previous studies conducted by Hirsch et al (Hirsch, Hayes et al. 2009), and contained three phases. Participants were required to undertake an initial 5-minute deep breathing focus period, followed by a 5-minute worry period, and then finally a second 5-minute deep breathing focus period.

During the pre- and post-breathing periods participants were instructed, using a standardized protocol, to focus their attention on their breathing, and if they find their concentration deviating, to redirect it back towards their breathing. Participants were not instructed to keep their eyes closed but were permitted to do so if they chose. During the worry period, participants were asked using a standardized script to identify a current worry, defined as *‘intrusive thoughts or images about potential future events or catastrophes that produce negative feelings when they occur’*. Participants then engaged in continuous worry about this topic for 5 minutes, according to instructions, as previously described (Hirsch, Hayes et al. 2009).

During the pre- and post-worry breathing periods, 12 beeps were presented. The inter-beep-interval ranged between 20 and 30s and a random list of intervals was generated to produce a standardized sequence of beeps that had varied intervals. On hearing the beep, participants reported if they were a) focused on their breathing (referred to as ‘breathing focuses’ henceforth) OR b) if their attention had diverted to a negative, positive or neutral thought (‘thought intrusions’). Using dedicated response sheets, the researcher documented the number of ‘breathing focuses’ and the valence (positive, negative, neutral) and content of thought intrusions.

The dependent measures for this task were 1) number of ‘breathing focuses’ pre- and post-worry (assessing maintenance of attention on task goal) and 2) number of negative, neutral and positive thought intrusions pre- and post-worry. Our primary hypotheses focused on the number of ‘breathing focuses’ as well as the number of negative and positive thought intrusions. We hypothesized a reduction in the number of ‘breathing focuses’ and positive thought intrusions, as well as an increase in negative thought intrusions, in the forced awakening group post-worry, compared with controls.

### Overnight declarative memory consolidation: word-pairs task

A word-pairs tasks was used to measure overnight declarative memory consolidation (Plihal and Born 1997). During encoding in the evening, participants were presented with 54 semantically related word-pairs on-screen for 4 seconds in a randomized order, with an inter-stimulus interval of one second. Participants were informed that recall would be tested immediately after the task and again the following morning. Four dummy pairs were added at the beginning and end of the list to control for primacy and recency effects (see supplementary table 1 for word list). Immediately following encoding, participants were presented with the first word of each of the pairs and required to type in the paired associate. Accuracy feedback was given followed by the correct word-pair displayed for 2 seconds to facilitate further encoding. In order to avoid ceiling effects, no learning criterion was enforced in order for the task to terminate (e.g., >60% accuracy level) as it was anticipated that our young healthy sample would exhibit good levels of performance. The following morning, participants were presented with the first word of the pair, in a newly randomized order, and asked to type the correctly paired associate, after which feedback and the correct word-pair was given. Number of correct word-pairs was recorded, and an overnight retention score was calculated by computing the difference in number of correct words retained between the immediate recall phase and the delayed recall phase, with greater score indicative of enhanced overnight memory consolidation.

### State questionnaires

Questionnaires were administered at eight hourly time-points throughout the day (08:00-15:00). The Karolinska Sleepiness Scale (Kaida, Takahashi et al. 2006) (KSS) [1 item, score 1– 9 (9 = maximum Sleepiness)] was used to measure subjective sleepiness. The Positive and Negative Affect Schedule (Crawford and Henry 2004) (PANAS) measured both positive and negative affect [10 positive and 10 negative items, scores on a 5-point likert scale with a possible range of 10-50]. The State Trait Anxiety Inventory (Spielberger 1970) (STAI) measured state anxiety [6 items, score 6-24 (24 = maximum anxiety)]. The State Difficulties in Emotion Regulation Scale (Lavender, Tull et al. 2015) (S-DERS) measured generalized state emotion regulation [21 items, scored 1-5, total score = 21-105 (105 = maximum emotion dysregulation)]. The KSS, STAI, and PANAS were administered hourly, whilst the S-DERS was administered every two hours. Mean daily scores were calculated per participant and entered into the analysis.

### Psychomotor vigilance task

Participants performed the psychomotor vigilance task (PVT) (Basner and Dinges 2011) at 11:20am post-sleep manipulation to assess vigilant attention. Participants were instructed to press the left mousepad button as quickly as possible when an asterisk (*) was presented on-screen. 110 trials were completed, and the inter-stimulus-interval (ISI) for each trial varied in duration between 1-10s. ISIs were selected from a randomized list of pre-defined ISIs ranging 1-10s, therefore the overall duration of the task remained consistent between participants (lasting approximately 12 minutes). Five practice trials preceded the experimental trials. The number of lapses (RTs > 500ms) were recorded and the mean reaction time for responses (including lapses and excluding responses <100ms), were calculated in accordance with previous guidelines (Basner and Dinges 2011) for each participant, with higher RTs indicating slower responses.

### Statistics and Analysis

#### Sample size

Our primary outcomes of interest were emotional perception, attention, and memory from the emotional test battery. Based on findings from previous between-group intervention studies utilizing the battery in healthy participants (Thomas, Dourish et al. 2014), we powered the study to detect moderate-to-large effects, should they exist. Assuming a minimum between-group effect size of 0.8, at 5% level of significance and 80% power, the required sample size was estimated at 50 participants (25 in each group).

#### Data handling and inspection

Prior to analysis, data cleaning was performed on the emotional test battery data to remove trials with RTs >3SDs from the mean or <200ms. Next, visual inspection was performed using histograms and box plots to assess the distribution of outcome data for all tasks. Skewness and Kurtosis values as well as statistical tests of normality (Shapiro Wilk’s Test), equality of variance (Levene’s Test), and sphericity (Mauchly’s Test) were used to interpret data assumptions. Several variables were non-normally distributed and were not corrected following square root or logarithmic transformations. Therefore, in the absence of a suitable non-parametric equivalent, testing proceeded using ANOVA, which is sufficiently robust to handle data with slight deviations from normality (Feir-Walsh and Toothaker 1974).

#### Statistical analyses

Primary data analysis was performed using ANOVA to test for between-group differences in emotion processing measures (dependent variables from the FERT, ECAT, EMEM, FDOT, EREC) and measures from the worry induction task (breathing focuses, negative intrusions, positive intrusions). Pre-to post change score on the word-pairs task and vigilance performance (RTs and number of lapses from the PVT) were analysed using independent *t*-tests. Normally distributed questionnaire data were analyzed using independent *t*-tests while non-normally distributed questionnaire data (STAI and S-DERS scores) were analyzed using non-parametric *t*-test equivalents (Mann-Whitney U test). Statistical significance was defined as *p* ≤0.05 in all instances. Cohen’s *d* effect sizes were calculated using mean differences and pooled standard deviations.

## Results

### Screening and baseline characteristics

5272 participants were assessed for eligibility using the online screening questionnaire, of which 516 provided complete screening questionnaires. 51 met criteria and took part in the study (38 females; mean age = 24 years, SD=3.64). A summary of baseline characteristics can be found in table 2.

**Table 2:** Participant demographics.

| Variable | Forced Awakening (FA) |  | Uninterrupted Sleep (US) |  |
| --- | --- | --- | --- | --- |
|  | Mean | SD | Mean | SD |
| N, count | 27 |  | 24 |  |
| Sex, count | 20F:7M |  | 18F:6M |  |
| Age, mean | 24.28 | 3.84 | 23.08 | 3.34 |
| PSQI, mean total | 2.20 | 1.18 | 1.71 | 1.04 |
| TST, *mins* (1-week actigraphy) | 421.22 | 43.13 | 426.31 | 43.02 |
| SE, % (1-week actigraphy) | 91.19 | 4.31 | 92.18 | 4.25 |
| TST, *mins* (1-week sleep diaries) | 420.86 | 50.23 | 421.34 | 55.64 |
| SE, % (1-week sleep diaries) | 89.65 | 5.63 | 88.17 | 5.03 |
| DERS, mean total | 61.68 | 11.47 | 58.21 | 9.59 |
| RRS, mean total | 35.12 | 11.45 | 31.75 | 8.61 |
| STAI, mean total | 38.96 | 5.35 | 38.38 | 5.68 |
| EPQ-RS, mean total | 1.52 | 1.69 | 1.54 | 1.86 |
| HADS-Anxiety, mean total | 2.75 | 2.16 | 2.64 | 2.06 |
| HADS-Depression, mean total | 0.90 | 1.11 | 1.10 | 1.23 |

Data are means (standard deviation) unless stated otherwise. Abbreviations: PSQI= Pittsburgh Sleep Quality Index, TST = Total Sleep Time, SEI = Sleep Efficiency, DERS = Difficulty in Emotion Regulation Scale, RRS= Ruminative Responses Scale, STAI = State Trait Anxiety Inventory, EQP-RS = Eysenck Personality Questionnaire – Revised Short Form, HADS= Hospital Anxiety and Depression Scale.

### Manipulation checks

#### Sleep

Consistent with the aims of the manipulation, participants in the experimental group had significantly greater WASO and significantly lower TST, time spent in N2, N3, and REM sleep stages (Cohen’s d for all ≥1.56). There were no group differences in N1 or SOL. Sleep variables for both groups are displayed in table 3. Participants in the uninterrupted sleep condition obtained on average 434 mins TST defined by PSG, consistent with normal habitual sleep times obtained during the baseline sleep monitoring phase 426mins (Actigraphy) and 412 (Sleep Diary) prior to the manipulation.

**Table 3:** Sleep variables by group.

| Variable | Forced Awakening (FA) |  | Uninterrupted Sleep (US) |  | *t* value | *p* value | Cohen's *d* |
| --- | --- | --- | --- | --- | --- | --- | --- |
|  | Mean | SD | Mean | SD |  |  |  |
| SOL, *mins* | 16.58 | 19.52 | 17.92 | 15.57 | -.253 | 0.80 | -0.08 |
| WASO, *mins* | 260.66 | 84.01 | 23.84 | 9.47 | -9.83 | <0.001 | 3.96 |
| TST *mins* | 229.98 | 83.74 | 434.57 | 39.04 | -10.80 | <0.001 | -3.14 |
| N1, *mins* | 45.63 | 26.99 | 35.65 | 45.53 | .96 | 0.34 | 0.27 |
| N2, *mins* | 98.59 | 57.90 | 216.25 | 45.69 | -7.88 | <0.001 | -2.26 |
| N3, *mins* | 47.87 | 26.37 | 97.37 | 36.19 | -5.58 | <0.001 | -1.56 |
| REM, *mins* | 30.76 | 27.69 | 83.39 | 28.49 | -6.60 | <0.001 | -1.87 |

Data are means (standard deviation) unless stated otherwise. Abbreviations: SOL = Sleep Onset Latency, WASO = Wake After Sleep Onset, TST = Total Sleep Time, SEI = Sleep Efficiency Index, SWS = Slow Wave Sleep, REM = Rapid Eye Movement, N1 = stage 1 sleep, N2 = stage 2 sleep, FA = Forced Awakening, US = Uninterrupted Sleep.

### Effects on emotion processing and task-related emotion regulation

#### Facial expression recognition task (FERT)

In terms of accuracy for identifying facial expressions there was no significant group x emotion interaction effect [*F* (6, 50) =.305, *p* =. 93]. Effect sizes for accuracy values for each type of expression ranged from *d* = -0.34 to 0.27 (See table 4). For reaction times there was similarly no significant emotion x group interaction effect [F (6, 50) = .974, *p* = .44], Effect sized ranged from *d* = -0.31 to 0.55. Further unplanned exploratory analyses were conducted on the number of misclassifications of each facial emotion, for which there was again no significant group x emotion interaction effect [*F* (6, 50) = .331, *p* =.92] (See supplementary table 2).

**Table 4:** Emotional Test Battery (ETB) values.

| Task | Emotion/valence | Group |  |  |  | Effect Size |  |  |
| --- | --- | --- | --- | --- | --- | --- | --- | --- |
| Facial Emotion Recognition Task (FERT) (Percentage accuracy) |  | Forced Awakening, |  | Uninterrupted Sleep, |  |  |  |  |
|  |  | Mean | SD | Mean | SD | Cohen's *d* | 95% CI |  |
|  | Anger | 62.96 | 6.20 | 60.52 | 11.08 | 0.27 | -0.27 | 0.82 |
|  | Disgust | 68.89 | 7.85 | 65.00 | 15.98 | 0.31 | -0.23 | 0.86 |
|  | Sadness | 67.41 | 7.02 | 65.63 | 10.08 | 0.21 | -0.34 | 0.75 |
|  | Fear | 54.72 | 16.16 | 54.06 | 16.15 | 0.04 | -0.50 | 0.58 |
|  | Surprise | 69.81 | 5.14 | 71.56 | 4.98 | -0.34 | -0.89 | 0.20 |
|  | Neutral | 81.11 | 12.50 | 79.58 | 17.81 | 0.10 | -0.44 | 0.64 |
|  | Happy | 80.46 | 5.42 | 79.79 | 6.42 | 0.11 | -0.43 | 0.66 |
| FERT (Reaction Times) | Anger | 1564.52 | 328.77 | 1497.12 | 226.58 | 0.24 | -0.33 | 0.80 |
|  | Disgust | 1578.85 | 442.58 | 1587.92 | 316.51 | -0.02 | -0.59 | 0.54 |
|  | Sadness | 1370.89 | 234.43 | 1332.71 | 197.30 | 0.18 | -0.39 | 0.74 |
|  | Fear | 1943.26 | 463.61 | 1857.79 | 348.66 | 0.21 | -0.36 | 0.77 |
|  | Surprise | 1501.59 | 361.43 | 1329.59 | 243.70 | 0.55 | -0.02 | 1.13 |
|  | Neutral | 1290.89 | 323.80 | 1391.04 | 314.68 | -0.31 | -0.88 | 0.25 |
|  | Happy | 1375.67 | 250.81 | 1266.17 | 198.64 | 0.48 | -0.09 | 1.05 |
| Faces Dot Probe Task (FDOT) (Vigilance Scores (ms)) | Fear (Masked) | -2.38 | 37.18 | 3.33 | 26.56 | -0.18 | -0.73 | 0.38 |
|  | Fear (Unmasked) | 2.58 | 28.61 | -.46 | 28.75 | 0.11 | -0.44 | 0.65 |
|  | Happy (Masked) | -7.96 | 55.81 | -3.75 | 32.19 | -0.09 | -0.64 | 0.45 |
|  | Happy (Unmasked) | 1.31 | 30.57 | -6.29 | 25.94 | 0.27 | -0.29 | 0.81 |
| Emotional Categorization Task (ECAT) (Reaction time (ms)) | Positive | 789.63 | 168.39 | 770.50 | 148.86 | 0.12 | -0.41 | 0.65 |
|  | Negative | 852.59 | 194.38 | 796.91 | 124.17 | 0.34 | -0.22 | 0.89 |
| Emotional Recall Task (EREC) (Number of correctly recalled words) | Positive | 4.67 | 3.17 | 5.08 | 2.57 | -0.14 | -0.69 | 0.40 |
|  | Negative | 3.59 | 2.96 | 4.29 | 2.39 | -0.26 | -0.81 | 0.29 |
| Emotional Recognition Memory Task (EMEM) (Percentage Accuracy) | Positive | 85.18 | 10.05 | 84.44 | 11.95 | 0.07 | -0.48 | 0.61 |
|  | Negative | 67.90 | 11.62 | 69.58 | 14.56 | -0.13 | -0.67 | 0.42 |

#### Faces dot-probe task (FDOT)

In terms of vigilance indices for masked faces there was no significant group x emotion interaction effect [*F* (1, 50) = .128, *p* = .72]. Similarly, there was no significant group x emotion interaction effect [*F* (1, 50) = .009, *p* = .93] for unmasked faces. Effect sizes ranged from *d* = -0.18 to 0.27 (see table 4).

#### Emotional categorization task (ECAT)

In terms of reaction times for positive and negative words, there was no significant valence x group interaction effect [*F* (6, 50) = .202, *p* = .65]. Effect sizes for positive, and negative words were *d* = 0.12 and 0.34 respectively (see table 4).

#### Emotional recall task (EREC)

In terms of the number of positive and negative words recalled, there was no significant group x valence interaction effect [*F* (1, 50) =.064, p = .80]. Effect sizes for positive and negative words were *d* = -0.14, and -0.26 respectively (see table 4).

#### Emotional recognition memory task (EMEM)

In terms of the number of positive and negative words recognised, there was no significant group x valence interaction effect [*F* (1, 50) = .256, *p* = .61]. Effect sizes for positive and negative words were *d* = 0.07 and -0.13 respectively (see table 4).

#### Worry induction task

We investigated ‘breathing focuses’ and positive/negative intrusions prior to and following the worry induction. In terms of the number of ‘breathing focuses’, there was no significant effect of group [*F* (1, 50) = .500, *p* = .483, *d* = 0.18] or time [*F* (1, 50) = .660, *p* = .420] and no significant group x time interaction effect [*F* (1, 50) = 1.630, *p* =.208]. For negative thought intrusions, there was no significant effect of group [*F* (1, 50) = 1.847, *p* =.180, *d* = 0.38] and no significant group x time interaction effect [*F* (1,50) =.034, *p* =.855]. However, there was a significant main effect of time [*F* (1,50) = 9.80, *p* = .003], with higher negative thought intrusions present in both groups post-worry, confirming the ability of the worry induction to induce negative intrusions. Similarly, there was no significant effect of group [*F* (1,50) = 0.72, *p* = 0.789, *d=* -0.44], or group x time interaction [*F* (1,50) = 1.647, *p* = .205] for positive thought intrusions, but there was a significant effect of time [*F* (1,50) = 6.14, *p* = 0.017] with fewer positive thought intrusions observed in both groups post-worry (see table 5).

**Table 5:** Breathing focus tasks data.

|  | Timepoint | Forced Awakening |  | Uninterrupted Sleep |  | Cohen's *d* | 95% CI |  | Time effect | Group*Time effect |
| --- | --- | --- | --- | --- | --- | --- | --- | --- | --- | --- |
|  |  | Mean | SD | Mean | SD |  |  |  | (*F*, *p*) | (*F*, *p*) |
| Breathing Focuses* | Pre-Worry | 8.11 | 1.97 | 8.83 | 2.08 | -0.36 | -0.90 | 0.20 | $F^{\dagger} = .660$ | $F^{\dagger} = 1.630$ |
| | Post-Worry | 8.22 | 2.21 | 8.33 | 2.78 | -0.04 | -0.90 | 0.59 | $p = .420$ | $p = .208$ |
|  | Pre-Worry | 8.22 | 2.21 | 8.33 | 2.78 | -0.04 | -0.90 | 0.59 |  |  |
| Negative Thought Intrusions | Pre-Worry | 1.00 | 1.33 | 0.50 | 0.98 | 0.42 | 0.14 | 0.97 | $F^{\dagger} = 9.80$ | $F^{\dagger} = .034$ |
| | Post-Worry | 1.44 | 1.74 | 1.00 | 1.18 | 0.29 | -0.26 | 0.84 | $p = .003$ | $p = .855$ |
| Positive Thought Intrusions | Pre-Worry | 1.26 | 1.16 | 1.54 | 1.56 | -0.21 | -0.21 | 0.75 | $F^{\dagger} = 6.14$ | $F^{\dagger} = 1.647$ |
| | Post-Worry | 1.07 | 1.14 | 0.96 | 1.04 | 0.10 | 0.10 | 0.45 | $p = .017$ | $p = .205$ |

Group means and variables. Data are mean (SD) \*number of times a participant-maintained focus on their breathing during timed breathing focus period. Time and group\*time effects for repeated measures ANOVA are presented <sup>†</sup>Degrees of freedom = 1, 50.

### Effects on vigilance and memory

#### Psychomotor vigilance task

Independent *t*-tests indicated significantly slower reaction times in the experimental group (mean RT= 428.79, SD= 57.59) compared with control [(mean RT= 390.03, SD=57.88), *p* = 0.023, *d* = 0.67] as well as increased attentional lapses (mean lapses = 7.63 SD=5.20 in experimental group vs. mean lapses = 4.12, SD=4.20 for the control group; *p* = 0.02, *d =* 0.74). See table 6.

#### Declarative memory consolidation

The number of correctly recalled word pairs at the evening time-point, pre-sleep, was similar between groups [experimental group [24.50 (SD 12.94)] vs. control group [25.43 (SD 10.12)].

An independent *t*-test of overnight retention scores demonstrated reduced overnight memory consolidation in the experimental group compared to the control group [overnight retention = 6.73 word-pairs (SD 4.13) for experimental group vs. 12.15 word-pairs (SD 5.79) for control group, *p* =.001, *d* = -1.08] (See table 6).

**Table 6:** Effects on vigilance, memory and self-report questionnaires.

|  | Forced Awakenings (FA) |  | Uninterrupted Sleep (US) |  |  |  | Cohen's *d* (95% CI) |  |  |
| --- | --- | --- | --- | --- | --- | --- | --- | --- | --- |
|  | Mean | SD | Mean | SD | t-value/u value* | *p*-value |  |  |  |
| PANAS +ve | 20.90 | 8.26 | 25.94 | 7.91 | 2.21 | 0.031 | -0.62 | -1.15 | -0.05 |
| PANAS -ve | 11.27 | 1.41 | 10.68 | 0.98 | -1.69 | 0.097 | 0.54 | -0.08 | 1.03 |
| STAI | 7.96 | 1.99 | 6.76 | 1.50 | -2.40* | 0.024 | 0.68 | 0.11 | 1.23 |
| S-DERS | 31.54 | 3.32 | 32.54 | 5.08 | 0.85* | 0.406 | -0.24 | -0.78 | 0.31 |
| KSS | 5.54 | 2.01 | 4.09 | 1.49 | -2.70 | 0.009 | 0.81 | 0.25 | 1.38 |
| PVT RT (*ms*) | 428.79 | 57.59 | 390.03 | 57.88 | 2.31 | 0.023 | 0.67 | 0.11 | 1.23 |
| PVT lapses | 7.63 | 5.20 | 4.12 | 4.20 | 2.65 | 0.015 | 0.74 | 0.18 | 1.30 |
| Word Pairs | 6.73 | 4.13 | 12.15 | 5.79 | 3.63 | 0.001 | -1.08 | -1.67 | -0.50 |
| **Memory task (Overnight improvement)** |  |  |  |  |  |  |  |  |  |

Data are mean (SD), effect sizes are *d* (95% CI) PANAS= Positive and Negative Affect Schedule, +ve = positive, -ve = negative, STAI = State Trait Anxiety Inventory, SDERS = State Difficulties with Emotion Regulation Scale, KSS=Karolinska Sleepiness Scale, PVT = Psychomotor Vigilance Task.

### Effects on self-reported affect, anxiety, sleepiness and emotion regulation

There were significantly higher levels of next-day state anxiety (STAI scores), sleepiness (KSS scores) and lower positive affect (PANAS) in the experimental vs. control group (medium-to-large effect sizes; see table 6). There was a medium effect size difference for negative affect (d=0.48), with the experimental group reporting increased scores, though this was not statistically significant (*p*=.097). No group effect was observed for emotion dysregulation (S-DERS Scores).

## Discussion

This study investigated the effects of sleep continuity disruption on emotion processing and regulation using a spectrum of measures with demonstrable sensitivity to depression (Harmer, Cowen et al. 2011). We hypothesized that sleep continuity disruption would: 1) engender negative biases across the domains of emotional perception, attention, and memory, and 2) impair emotion regulation in response to worry induction. However, no significant group effects were observed, despite impairments in vigilant attention, overnight memory consolidation, and self-reported affect and anxiety.

Consistent with our manipulation aims, we observed large increases in WASO, alongside marked reduction in total sleep time, N3, REM, and psychomotor vigilance in the experimental group vs. control. Self-report data further indicated that participants in the experimental group were significantly sleepier, and had increased anxiety and reduced positive affect with moderate effect sizes. Finally, a large group effect was observed for overnight memory consolidation, with participants in the experimental group exhibiting poorer overnight retention of (non-emotional) word-pairs. Together these results suggest that sleep discontinuity had a significant impact on daytime functioning measures known to be sensitive to sleep disruption (Plihal and Born 1997, Finan, Quartana et al. 2017).

Tests of difference in emotion processing variables yielded no significant findings for facial emotion recognition, categorization, recall or recognition of emotional words, nor attentional biases towards emotional faces. While difficult to directly compare with previous studies due to their small sample sizes and paradigm differences, our findings are largely contrary to historical reports indicating large effects of sleep disruption on emotion processing, particularly with regards to facial emotion processing (Van Der Helm, Gujar et al. 2010). Nevertheless, a recent meta-analysis (Tomaso, Johnson et al. 2020) found only small effects of sleep deprivation on emotion processing (g = -0.11) and adaptive emotion regulation (g=-0.32), whereas effects of self-report positive (g = -0.94) and negative mood (g=0.45) were greater indicating the effects of insufficient sleep on objective measures of emotion might not be as strong as previously believed, consistent with the significant (subjective mood) and non-significant effect sizes (emotion processing) we report. Moreover, recent large-scale studies (n=291 (Holding, Laukka et al. 2017), n=201(Brand, Schilling et al. 2019)) also showed no effect of sleep disruption (Holding, Laukka et al. 2017, Gerhardsson, Åkerstedt et al. 2019, Tamm, Schwarz et al. 2020), total sleep time (Holding, Laukka et al. 2017), subjective sleep quality (Holding, Laukka et al. 2017), nor insomnia symptom severity (Brand, Schilling et al. 2019) (ISI scores) on emotional processing. Our findings suggest that experimentally induced sleep fragmentation over one night also has limited effect on emotional processing.

For the worry induction task, we found a main effect of time, reflecting an increase in negative and a decrease in positive thought intrusions (supporting task validity), yet no group effect of sleep disruption. While further examination of this task in the context of sleep is necessary, our findings suggest that one night of sleep fragmentation does not influence the ability to regulate intrusive thoughts under emotion challenge (worry). Supporting our findings, recent work also confirmed the absence of a causal effect of sleep deprivation on emotion regulation, demonstrating no effect on a cognitive re-appraisal task following 24h of total sleep deprivation (Shermohammed, Kordyban et al. 2020).

Nevertheless, our findings should be considered in the context of the strengths and limitations of the study design. We disrupted sleep continuity for just one night; multiple nights of sleep disruption may be required to create negative bias in emotional processing. Furthermore, our study does not isolate a number of factors known to be of clinical relevance for insomnia disorder, such as perception of and beliefs about sleep quality. Continued development of sleep manipulation paradigms which tap into these clinical constructs is required to translate these findings into insomnia populations. Second, our sample was predominately female (74.5%), reasonably small, a between-subject design was used, and the study was only powered to detect effects anticipated within the medium to large range. Effect sizes for group differences on the emotional tasks were chiefly in the range of *d*=0.1-0.3, which aligns with recent meta-analytic data (Tomaso, Johnson et al. 2020) demonstrating an effect size of *g* = 0.1 on emotion processing. However, there was no consistency to the direction of these effects across variables. For example, although effects in the d=0.1-0.2 range were observed for RT to fearful faces, and accuracy for happy faces, yet in the opposite direction than hypothesized, which does not support the argument that effects were simply too discrete to be detected. Furthermore, this raises methodological issues as detection of such discrete effects would require sample sizes between n=350-1200 in each arm (assuming a between-subjects design where *d* = 0.1-0.3 and one-tailed power of 80%). Our study also has several strengths. Objective measurements of sleep were obtained using PSG, and participants were rigorously screened. Timing of the key emotional variables (emotional test battery) was standardized. Central hypotheses surrounding emotion processing and emotion regulation were tested across various domains and emotional constructs using valid probes with documented sensitivity to depression pathophysiology and manipulations in healthy volunteers.

To conclude, the present study observed that one night of sleep continuity disruption, using an experimental model of insomnia, had statistically significant cognitive and affective consequences, but was not associated with comparable effects across multiple domains of emotion processing, despite impairments in declarative memory consolidation, vigilance, positive affect and anxiety. This study therefore suggests that, although one night of sleep continuity disruption has clear cognitive and affective consequences, it does not impair emotion processing or regulation. Such findings may develop existing knowledge of how sleep disruption leverages mechanistic processes that promote risk for depression. Future studies may wish to utilize within-subject paradigms that may address effect size considerations by optimising statistical power within obtainable sample sizes, however caution should be exercised to ensure suitability of outcome measures for repeated measures designs. Future studies may also wish to extend this work by considering the use of alterative populations, such as those with underlying personality traits (e.g. neuroticism), or habitually poor sleep, which may confer a greater risk profile for emotional pertubation, consistent with stress-diathesis models (Slavik and Croake 2006) of mental illness. Finally, our work may be extended by probing the potential role that respective sleep stages may play in mediation of both mood and emotion processing responses.

## Conflict of Interest Statement

The authors of this manuscript declare no financial, or non-financial conflicts of interest

## Disclosure Statement

Financial Disclosure: none. Non-financial Disclosure: none This research study was supported financially by the National Institute for Health Research (NIHR) Oxford Biomedical Research Centre (BRC). The research was also supported through a DPhil Scholarship (to M.J.R) from the Dr Mortimer and Theresa Sackler Foundation.

## Emotional Test Battery Methods

The Emotional Test Battery was delivered by administering the following tasks in a standardized fixed order:

### Facial emotion recognition: facial emotion recognition test (FERT)

In the facial emotion recognition test (FERT) participants categorize faces (500ms presentation time) with seven different facial expressions (happiness, sadness, fear, anger, disgust, surprise or neutral) displayed at morphed intensity levels from 0% (neutral) to 100% (full expression) (6 emotions × 10 intensities × 4 examples = 240 trials)^50^. Our a priori hypotheses were that the experimental group, relative to control, would perform with higher percentage accuracy and shorter reaction times for anger, disgust, sadness, fear, as well as lower accuracy and longer reaction times for surprise and happiness. Further unplanned analyses were also conducted on the number of times a facial stimulus was incorrectly classified as an emotion (misclassifications).

### Perception of emotional words: Emotion categorisation task (ECAT)

The emotion categorisation task (ECAT)^47,49^ assesses the speed of response towards positive and negative self-referent personality descriptors. Sixty personality characteristics selected to be disagreeable (e.g. “domineering”) or agreeable (e.g. “cheerful”), matched for frequency, length, and meaningfulness were presented for 500ms in a random order. Participants then expressed whether they would ‘like’ or ‘dislike’ to be referred to by each characteristic word. participants were not instructed to memorize the words. The outcome measure was reaction time for correct responses to positive and negative words. We hypothesized shorter reaction times for negative words, and longer reaction times for positive words in the experimental vs control group.

### Attentional bias for emotional faces: faces dot probe task (FDOT)

The faces dot probe task^51^ (FDOT) provides an assessment of relative attention to positive and negative emotional faces. Following a fixation cross, two faces were simultaneously presented on the computer screen, one above the other. Faces displayed either positive (happy), negative (fear) or neutral facial expressions (either a positive-neutral pair or a negative-neutral pair) and were then replaced by a pair of white dots (:), which appeared behind the top or the bottom face, with equal frequency. Probes appeared behind the emotional (congruent trials) or neutral (incongruent trials) faces. Participants then responded by indicating the orientation of the probe. 96 trials were completed (48 happiness, 48 fear). On 50% of trials, faces were presented for 500ms (unmasked). On the remaining 50% of trials, the faces were presented for 14ms followed immediately by a mask consisting of special characters (masked). The dependent variables were relative vigilance scores^a^ (: (calculated as *mean RT emotion-congruent trials* – *mean RT emotion-incongruent trials*, measured in milliseconds)) for each emotion (masked and unmasked). Higher vigilance score reflected a greater attentional bias towards the relevant emotional valence and thus we hypothesized higher vigilance indices for fearful faces and lower vigilance indices for happy faces in the experimental vs control group.

### Emotional recall memory: emotional recall task (EREC)

Participants were asked to write down (on paper) as many of the words previously presented in the emotional categorisation task (ECAT), in a 2-minute interval. Participants were not informed of this task prior to commencing the ETB, and thus was a surprise recall. The recall of emotional words gives a measure of bias for both positive and negative words. The outcome measure was the number of positive and negative self-referent words correctly recalled, thus we hypothesized higher recall of negative words and lower recall of positive words in the experimental vs control group.

### Emotional recognition memory: emotional recognition memory Task (EMEM)

Participants were presented with 60 positive and 60 negative self-referent personality descriptors for 500ms in a random order. Some were previously seen in the ECAT, some were unseen distracter words. Participants reported if they had previously seen the word during the ECAT. Outcome measures were the percentage of correct responses for positive and negative words. We hypothesized lower recognition accuracy for positive words and higher recognition accuracy for negative words in the experimental vs control group.

## Detailed Hypotheses

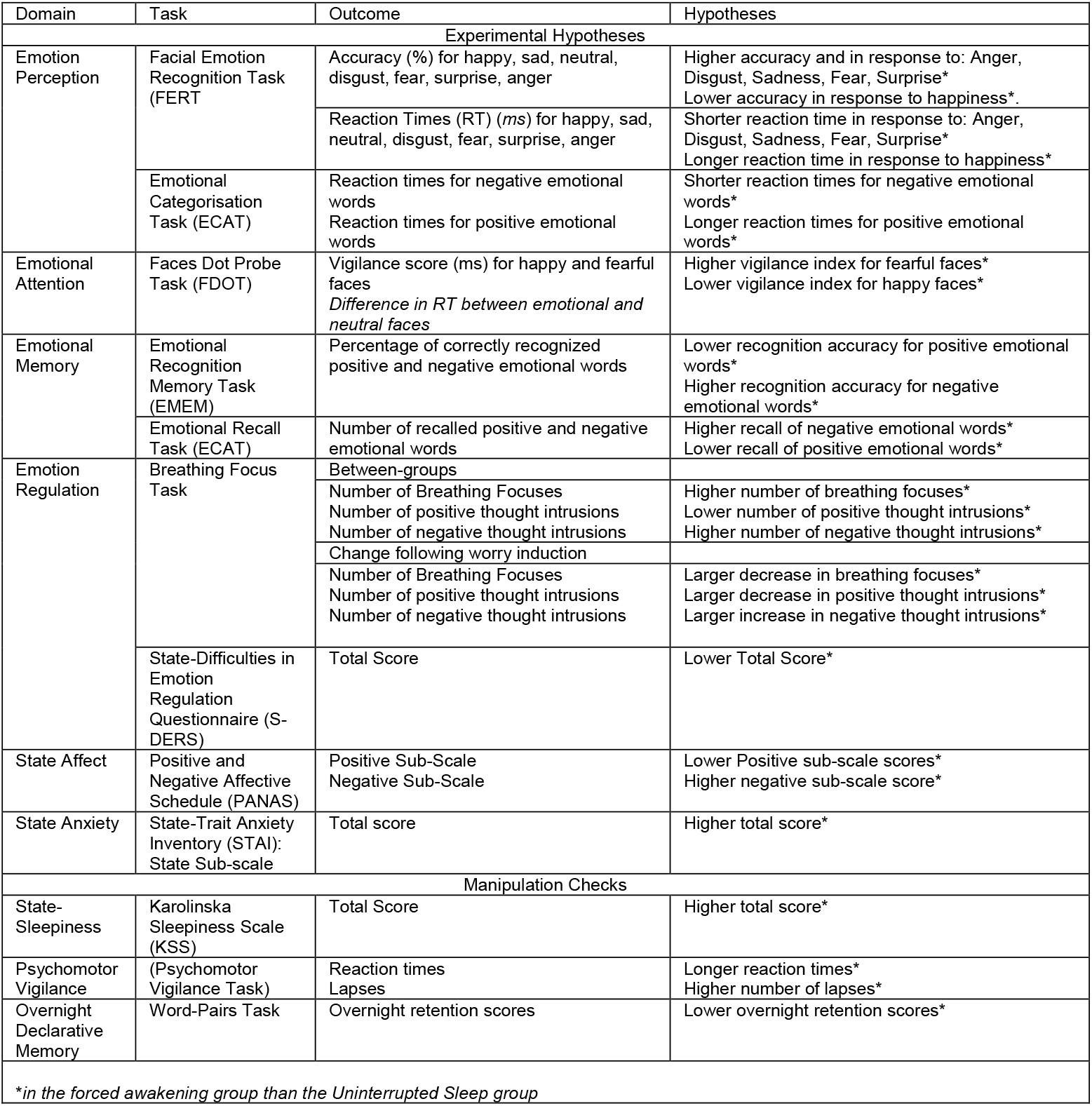

